# In-cell Proteomics Enables High-Resolution Spatial and Temporal Mapping of Early *Xenopus tropicalis* Embryos

**DOI:** 10.1101/2025.05.23.655823

**Authors:** Jian Sun, Xiaolu Xu, Shuo Wei, Yanbao Yu

**Affiliations:** Department of Biological Sciences, University of Delaware, 105 The Green, Newark, Delaware, 19716 (USA); Department of Chemistry and Biochemistry, University of Delaware, 163 The Green, Newark, Delaware, 19716 (USA)

**Keywords:** animal–vegetal axis, E4technology, on-filter in-cell (OFIC) digestion, in-cell proteomics, Spatiotemporal proteomics, *Xenopus tropicalis*

## Abstract

Early embryonic development requires tightly regulated molecular programs to coordinate cell division, fate specification, and spatial patterning. While transcriptomic profiling is widely performed, proteomic analyses of early vertebrate embryos remain limited due to technical challenges in embryonic sample preparation. Here, we propose an “in-cell proteomics” approach, which bypasses cell lysis and yolk depletion, processes individual embryos directly in functionalized filter devices, and generates liquid chromatography–mass spectrometry (LC-MS)-friendly samples in an extremely robust and streamlined manner. Combined with a single-shot data-independent acquisition (DIA) MS workflow, this approach enabled us to consistently quantify ∼6,200 proteins from a single *Xenopus tropicalis* embryo, representing the deepest proteomic coverage of early *X. tropicalis* development reported to date. Investigation of the temporal proteomes across five cleavage stages (1- to 16-cell) revealed a drastic proteomic shift between 2- and 4-cell stages, followed by more gradual transitions thereafter. Spatial analysis of dissected 8-cell blastomeres uncovered pronounced molecular asymmetry along the animal–vegetal axis, while dorsal–ventral differences were minimal. This study establishes a novel in-cell proteomics technology in conjunction with DIA-MS as a robust platform for high-resolution, low-input developmental proteomics analysis, and provides a comprehensive spatiotemporal protein atlas for early *X. tropicalis* embryos.

---

Early vertebrate embryogenesis is driven by a series of precisely orchestrated molecular and cellular events that progressively specify cell fates and establish the body plan.^[1, 2]^ In amphibians such as *Xenopus*, as in other vertebrates, this process begins with rapid, synchronous cleavage divisions that generate blastomeres with distinct developmental potentials along the animal–vegetal and dorsal–ventral axes.^[3-5]^ *Xenopus tropicalis*, a relative of the more widely used allotetraploid *X. laevis*, is smaller—about one-third the body size and one-fifth the embryonic material.^[6]^ Due to its shorter generation time and true diploid genome, *X. tropicalis* has emerged as an excellent model for early vertebrate development.^[7, 8]^ While transcriptional regulation during early embryogenesis has been extensively studied,[9] in particular using *X. laevis* embryos, the proteomic landscape of *X. tropicalis* remains largely unexplored, mainly due to challenges by the minute amounts of starting material and technical difficulties in sample preparation. For instance, the presence of abundant yolk proteins in every embryonic cell has severely hindered the detection of low-abundance proteins in *Xenopus* embryos.^[10-12]^ Although new yolk depletion methods have been developed to improve MS detection, these procedures are time-consuming and labor-intensive.^[10]^ Therefore, there are urgent unmet needs to develop effective and robust ways to examine the proteomes of *X. tropicalis* embryos at various developmental stages with low or no technical barriers.

Recent innovations in low-input proteomics, particularly the on-filter in-cell (OFIC) processing based E4technology, have begun to overcome these limitations.^[13, 14]^ This single-vessel format technology enables direct in situ protein digestion within methanol-fixed cells, the so-called “in-cell proteomics”, thus greatly reducing sample handling and loss, and simplifying the proteomics workflow. This method has been shown to achieve unbiased and in-depth proteome coverage in yeast, mammalian cells, and *Caenorhabditis elegans*.^[13, 14]^ Importantly, the E4technology is compatible with low-cell or low-input applications, offering a practical solution for developmental studies where starting material is inherently limited. In the present study, we leveraged OFIC processing to analyze the temporal and spatial proteome dynamics of early *X. tropicalis* embryos. Specifically, we investigated spatiotemporal proteomes spanning five cleavage stages (one-cell to sixteen-cell) and spatially distinct blastomeres at the 8-cell stage.

Our first experiment was to investigate if the in-cell proteomics is applicable to embryonic samples. We collected individual *X. tropicalis* embryos and loaded directly to the filter device, E4tips, and then fixed the whole embryo with methanol. The fixed embryos were then subjected to tryptic digestion after reduction and alkylation in the cells. With no fractionation or yolk depletion, we were able to identify over 5,300 proteins from a single *X. tropicalis* embryo. Compared with the conventional SDS-based lysis method, the in-cell digestion not only yielded consistently higher number of proteins and peptides identified, but also greatly reduced variabilities (**Figure S1**). Encouraged by these initial results, we set out to further investigate how we can leverage this method to answer biologically significant questions.

We then analyzed the temporal proteome of the early cleavage stages, from 1-to 16-cell, with four biological replicates per stage (**Figure 1**). The in-cell proteomics approach enabled consistent identification from each one of the 20 LCMS runs (**Figure 2A**), resulting in a total of 6,375 non-redundant protein groups. As proteomics investigations of X. tropicalis embryos are rarely seen, to our best knowledge, this study represents the largest *X. tropicalis* embryonic proteome reported to date (**Table S1**). Among them, 6,277 proteins were detected at all five stages (**Figure 2E**), suggesting large qualitative similarities among the early embryonic proteomes. Based on the Xenbase annotation,^[15]^ the protein identified also included 331 transcription factors (**Figure 2H**), which are known to play crucial roles in early development but tend to have relatively low abundance. The frog embryo proteome spanned over six orders of magnitude (**Figure S2A**). The three yolk precursor proteins-Vitellogenin-B2, A2, and A1-ranked top3 among the entire proteome and accounted for 42% of the total protein mass. The top15 most abundant proteins contributed to over half of the embryonic proteome (**Figure 2H**). These data underscore the high dynamic range of embryonic proteome and the extreme challenge for low-abundance protein analysis. Previously, it was believed that the yolk proteins can make up >90% of the total proteome in the embryos, and yolk depletion has been exercised to enhance proteome coverage.^[10]^ Our findings suggested that the in-cell proteomics approach and the single-run DIA LC-MS can provide unprecedented proteomic depth, eliminating the need for yolk depletion or peptide pre-fractionation.

**Figure 1.**
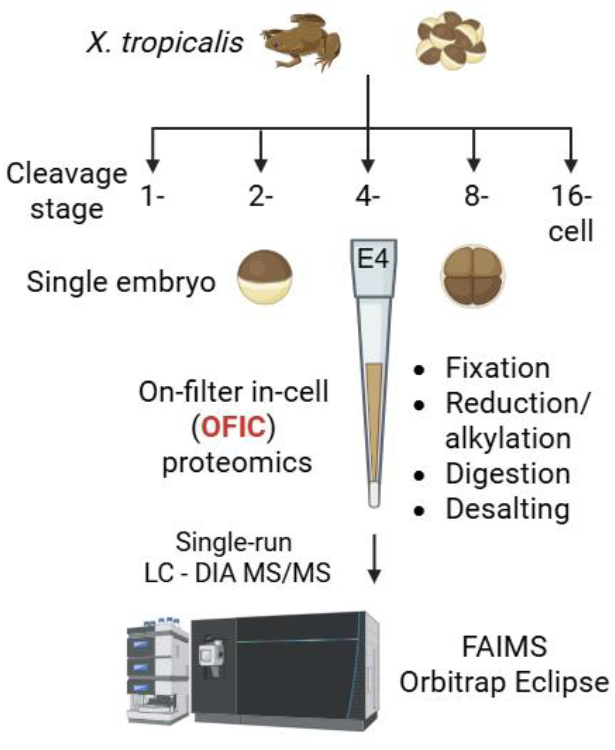
Workflow of the “in-cell proteomics” approach for *X. tropicalis* embryos analysis. Embryos were collected at 1-, 2-, 4-, 8-, and 16-cell stage, respectively. One embryo was used for each digestion experiment, and four embryos (biological replicates) were processed independently for each cleavage stage using E4tip. Mass spectrometric analysis was performed using Orbitrap Eclipse with FAIMS Pro Interface in DIA mode.

**Figure 2.**
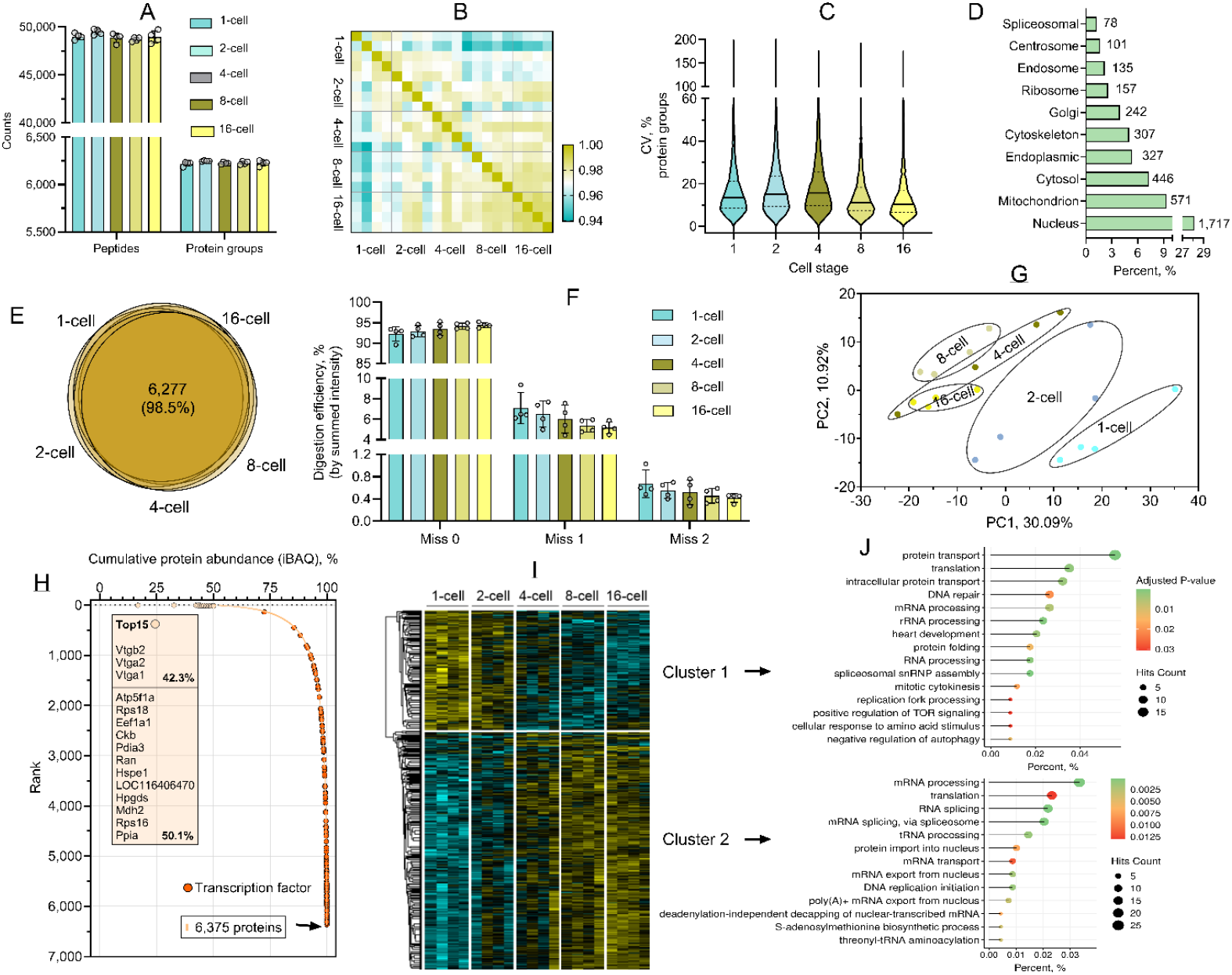
Evaluation of the in-cell proteomics approach for *X. tropicalis* embryo proteomic analysis. (A) Identification of protein groups and peptides across the five cleavage stages. Error bars indicate four biological replicates. (B) Pearson correlation analysis. (C) Coefficient of variation of the protein groups among the five stages. Horizontal solid lines indicate the median value. (D) Cellular compartment analysis of the overall *X. tropicalis* embryo proteome. (E) Venn diagrams. The number and percentage indicate the shared proteins among the five stages. (F) Digestion efficiency. The summed intensity of peptides carrying one and two missed cleavages were divided by the total peptide intensity. (G) Principal component analysis. (H) Protein rank plot. The top15 most abundant proteins were indicated in the text box. Transcription factors are highlighted in orange. (I) Heatmap of the ANOVA significant (FDR 0.05) proteins among the five stages. Two clusters were highlighted. (J) Gene ontology biological process analysis of proteins in Cluster 1 and Cluster 2, respectively.

In terms of technical performance of the OFIC approach, among the biological replicates at each stage, the Pearson correlation averaged from 0.98 to 0.99, and the median of the protein group coefficient of variation (CV) ranged from 10-16% (**Figures 2, B-C**), indicating excellent reproducibility. On the other hand, as the enzymatic penetration and in-cell digestion might be restricted by the embryo texture, we assessed the trypsin digestion efficiency by examining the number of peptides carrying un-cleaved sites as well as their intensity (**Figures 2F** and **Figure S2B**). Our data implicate that only around 22% of the total number of peptides contained one or two miscleavages, which account for 8% or less of the total peptide intensity, similar to lysate-based digestion experiments reported before.^[13]^ These data suggest that in-cell digestion of Xenopus embryos after a simple methanol fixation is entirely feasible.

Subcellular localization analysis (**Figure 2D**) revealed a broad representation of cellular compartments, with the majority of proteins mapped to the nucleus (28%), mitochondrion (9.4%), and cytosol (7.3%). Additional enrichment was observed in the endoplasmic reticulum, ribosome, Golgi apparatus, and lysosome, indicating comprehensive proteome coverage across diverse cellular compartments. When compared with *X. laevis* data derived from a lysate-based digestion method,^[16]^ the proteins derived from OFIC approach did not show significant bias toward any particular cellular localization (**Figure S3)**. This finding is consistent with the data from in-cell digestion of *C. elegans* worms reported recently.^[14]^

Next, we examined the proteome-wide differences among the five developmental stages. A simple principal component analysis (PCA) using the global proteome profiles without imputation can already classify the cleavages (**Figure 2G**). While there were partial overlaps among 4-, 8-, and 16-cell-stage embryos, a clear segregation can be seen between the 2- and 4-cell stages, indicating a major proteomic transition during the early developmental window and a more gradual trajectory of proteomic changes at later cleavage stages. Multi-sample test (ANOVA, Permutation FDR 0.05) corroborated the findings, and revealed over 1,000 differential proteins and two distinct clusters between stages of 1- and 2-cell (Cluster 1) and 4/8/16-cell (Cluster 2) (**Figure 2I**), highlighting a key transition in the temporal progression of the early embryo proteome. The 363 proteins upregulated at earlier stages (Cluster 1) were enriched in pathways related to protein metabolism, such as protein translation, folding, and transport, likely reflecting the earliest embryos’ reliance on maternal proteins and proteins translated from maternal mRNAs prior to the maternal to zygotic transition. Conversely, the 703 proteins upregulated at later stages (Cluster 2) were enriched in pathways related to RNA metabolism, including RNA processing, transport, and translation, suggesting the activation of RNA regulatory pathways to support zygotic genome activation (**Figure 2J**). While there was some functional overlap between the two clusters, such as enrichment in translation and DNA replication, these categories likely reflect distinct subsets of proteins undergoing turnover versus production. Taken together, the temporal proteomic profiles highlight a tightly regulated, stage-specific remodeling of the embryonic proteome, with the 2- to 4-cell transition representing a major developmental inflection point marked by dynamic proteomic reprogramming and fine-tuned regulation of core cellular processes.

We further leveraged the E4technology to examine the individually dissected blastomeres from 8-cell stage embryos (**Figure 3A** and **Table S2**). The four blastomeres—D1 (dorsal-animal; future central nervous system), V1 (ventral-animal; future neural crest and epidermis), D2 (dorsal-vegetal; future trunk mesoderm), and V2 (ventral-vegetal; future trunk endoderm)^[17]^—were obtained with five biological replicates per lineage (1 blastomere per embryo; five embryos). Similar to whole embryo samples, the blastomeres were processed entirely in the E4tips (**Figure 3A**). The streamlined workflow once again offered consistent identification rate (**Figure 3B**), 5,000-6,000 protein hits from each single blastomere, and minimal variability, as demonstrated by low CV and high reproducibility (**Figure 3, B-C**). Unsupervised hierarchical clustering analysis revealed distinct proteomic patterns between animal pole (D1 and V1) and vegetal pole (D2 and V2) blastomeres (**Figure S4**), highlighting spatially organized proteomic signatures. The PCA further supported this separation, with PC1 (45.6%) capturing the animal–vegetal distinction and PC2 (10.4%) distinguishing between D1 vs. V1 and D2 vs. V2 (**Figure 3E**). These findings indicate that spatially distinct proteomic programs are already established at the 8-cell stage. ANOVA test (Benjamini-Hochberg FDR 0.01) reported 2,024 significantly differential proteins and two major molecular asymmetries across the animal-vegetal axis (**Figure 3G** and **Table S2**). Cluster I encompassed 1,277 proteins enriched in animal pole blastomeres (D1 and V1), which were significantly enriched in pathways associated with anabolic processes such as DNA, RNA and protein synthesis, processing and quality control (**Figure 3H**). In contrast, Cluster II comprised 747 proteins upregulated in vegetal blastomeres (D2 and V2), which showed upregulation of pathways related to catabolic processes—including necroptosis, autophagy, endocytosis, lysosome, and peroxisome (**Figure 3H**). These signatures are consistent with the known divergence between the two sides, with the animal pole displaying more proliferative activity and the vegetal pole providing the energy and building blocks for growth.^[18]^ Interestingly, a direct comparison between dorsal (D1 + D2) and ventral (V1 + V2) blastomeres revealed minimal differences (**Figure 3F**). Only one protein—HNRNPA1, involved in mRNA processing^[19]^—was significantly upregulated (FDR < 0.01) in dorsal cells. Taken together, these spatially distinct proteomic signatures underscore the early emergence of lineage-specific molecular programs along the animal–vegetal axis of the 8-cell embryo, and suggests that dorsal–ventral proteomic asymmetry is limited at this stage, in contrast to the pronounced animal–vegetal divergence.

**Figure 3.**
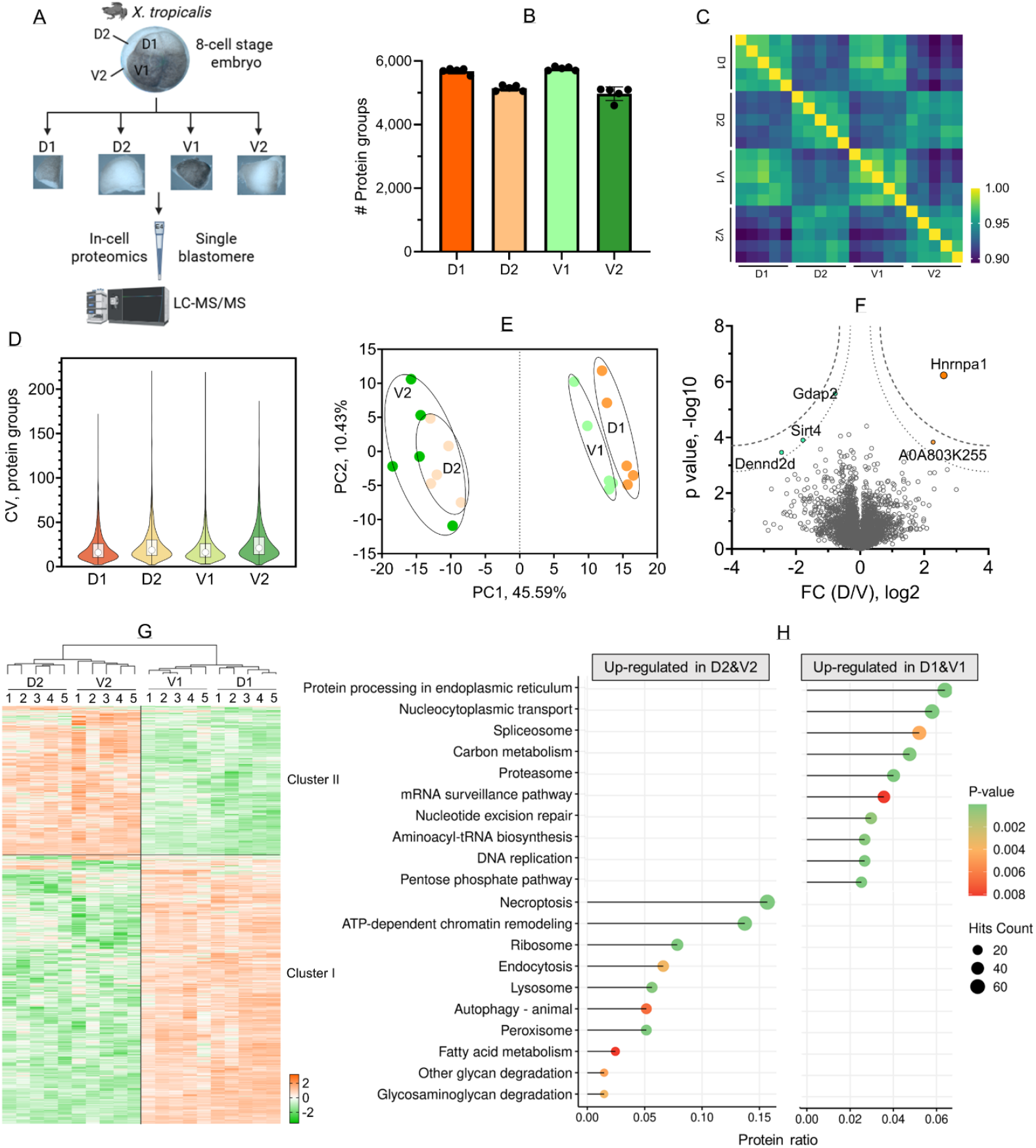
Spatial proteomic profiling of 8-cell stage *X*. tropicalis embryos using the OFIC approach. (A) Workflow of the OFIC approach for spatial proteomics analysis of 8-cell stage *X. tropicalis* embryos (B) Protein group identifications. (C) Pearson correlation. (D) Protein groups CV. (E) Principal component analysis. (F) Volcano plot of comparison between dorsal and ventral blastomeres. Dotted lines indicate Permutation FDR 0.05 and 0.01, respectively. (G) Heatmap of ANOVA significant proteins (FDR 0.01). (H) KEGG pathway analysis of the two cluster of proteins shown in panel G. Top10 enriched terms of each cluster were plotted.

In this study, we establish an “in-cell proteomics” platform as a robust and sensitive approach for developmental proteomics in early *X. tropicalis* embryos. This platform enables in-depth, reproducible protein quantification directly from single embryos and individual blastomeres without the need for yolk depletion or peptide fractionation, overcoming long-standing technical barriers in low-input embryonic proteomics. The strategy of in-cell processing preserves sample integrity and minimizes protein loss, allowing consistent detection of over 6,000 proteins per sample, including low-abundance transcription factors. As global proteomics studies of *X. tropicalis* embryos are rare, this dataset represents the deepest proteomic coverage reported to date for early *Xenopus* development. The method’s high reproducibility and minimal variability highlight its suitability for quantitative proteomics even in highly yolk-rich embryonic systems.

Temporally, our analysis uncovered a sharp and well-defined proteomic transition between 2- and 4-cell stages, which likely marks a key developmental inflection point. This shift may reflect the onset of zygotic influence over cellular programs, as the embryo transitions from reliance on maternally deposited mRNA and proteins to active biosynthesis and regulatory control.^[5, 20, 21]^ Although zygotic genome activation in *Xenopus* typically occurs at the mid-blastula transition (MBT), earlier remodeling at the proteomic level likely prepares cells for rapid cleavage and downstream patterning. The distinct clustering of developmental stages further supports that proteomic reprogramming is not continuous but involves discrete transitions during early embryogenesis.

Spatially, proteomic profiling of individual blastomeres from 8-cell embryos revealed pronounced molecular asymmetries along the animal–vegetal axis. Animal pole blastomeres exhibited signatures of elevated biosynthesis, which is consistent with ectodermal lineage priming. In contrast, vegetal blastomeres displayed enriched pathways in catabolic processes such as lysosome and peroxisome function, reflecting early mesendodermal programming. These results provide proteomic evidence for molecular specialization of blastomeres well before morphological differentiation becomes evident.

In contrast, dorsal–ventral differences were minimal at the proteomic level. This observation aligns with the concept that dorsal–ventral axis formation is driven more by localized signaling cues, such as Wnt and Nodal pathway activators, and post-translational regulation, rather than differences in overall protein abundance at this early stage.^[22, 23]^ Furthermore, such determinants may be present at low copy numbers or sequestered in subcellular domains, remaining below detection thresholds in bulk proteomic analysis. Alternatively, their asymmetric activity may depend on dynamic protein modifications or RNA-level localization, neither of which was captured in this dataset.

Together, our findings establish E4technology with DIA-MS as a powerful and scalable strategy for high-resolution developmental proteomics, particularly well-suited for systems with limited input material and high yolk content. The spatiotemporal proteomic atlas provided here offers valuable insight into early embryonic patterning and lays the foundation for future investigations into lineage specification and molecular asymmetry during vertebrate development.

## Acknowledgements

We would like to acknowledge support from the National Institute of General Medical Sciences (NIGMS) under award number P20GM104316 for the Orbitrap Eclipse MS instrument. This work is also supported by NIH R01 DE029802 (S. W.). The table of content figure and some of the main figures were created with BioRender.

## Table of Contents

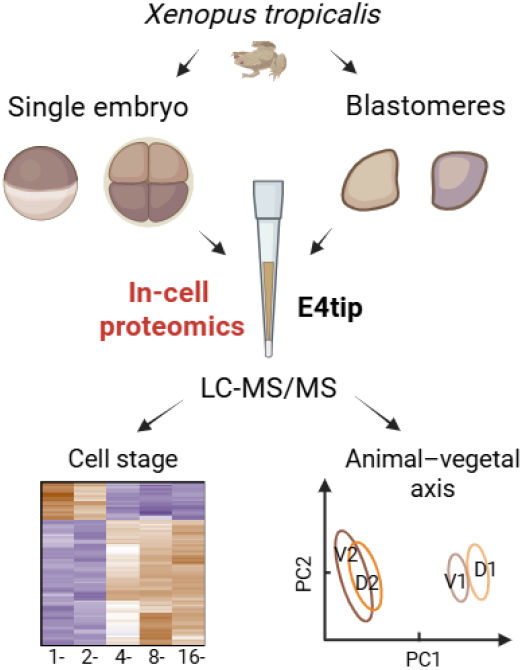

### Text for Table of Contents

A streamlined in-cell proteomics approach was developed for the first time for spatiotemporal characterization of *Xenopus* embryos. The approach eliminates cell lysis, and allows intact embryos to be processed into purified peptides in a single device with unprecedent proteome coverage. The great simplicity of the approach, in combination with single-run DIA MS/MS, offers a robust proteomics platform for studying developmental biology, and beyond.

