## Supplementary methods and figures for "In-cell Proteomics Enables High-Resolution Spatial and Temporal Mapping of Early *Xenopus tropicalis* Embryos"

### MATERIALS AND METHODS

#### *Xenopus tropicalis* embryo culture and collection

Wild-type *X. tropicalis* frogs were purchased from National *Xenopus* Resource in Woods Hole, Massachusetts. Methods involving live animals were carried out in accordance with the guidelines and regulations approved and enforced by the Institutional Animal Care and Use Committees at University of Delaware. *Xenopus tropicalis* embryos were generated using standard fertilization and culturing protocols.[1] For temporal proteomic analysis, embryos at five developmental stages—1-cell, 2-cell, 4-cell, 8-cell, and 16-cell—were collected. Four embryos (biological replicates) per stage were collected. For spatial proteomic analysis, 20 embryos at the 8-cell stage were harvested and fixed in 100% methanol. Embryos were then manually dissected into four lineage-representative blastomeres: D1 (dorsal-animal), V1 (ventral-animal), D2 (dorsalvegetal), and V2 (ventral-vegetal). For each blastomere group, five individually dissected cells were pooled to form one biological replicate, resulting in five replicates per group.

#### Protein sample preparation

For in-cell sample preparation, individual embryos or blastomeres were collected and transferred to E4tips XL (CDS Analytical, Oxford, PA) that were prefilled with 200 μl of methanol, and incubated on ice for 30 minutes. The tips containing fixed embryos were spun at 1,500 x g for one minute. The filters were washed one time with 200 μl of methanol. Then 100 μl of 50 mM triethylammonium bicarbonate (TEAB) with final concentration of 10 mM Tris (2-carboxyethyl) phosphine (TCEP) and 40 mM chloroacetamide (CAA) was added, followed by incubation at 45ºC for 10-15 min. The E4tips were then washed one time with 200 μl 50 mM TEAB. For protein digestion, 150 μl 50 mM TEAB plus 1.0 μg of trypsin/Lys-C mix (Promega, WI) was added to the samples and incubated at 37ºC for 16-18 hours with gentle shaking (350 rpm/min). After digestion, the samples were acidified with 1% formic acid (final concentration) and spun at low speed (400-600 x g) for 10 min. The tips were washed one time with 200 μl of wash buffer (0.5% acetic acid in water) to discard flow through. The tips were transferred to new collection tubes, and subjected to two sequential elution with 200 μl of Elution buffer I (60% acetonitrile and 0.5% acetic acid in water) and Elution buffer II (80% ACN and 0.5% acetic acid in water), respectively. The elution was pooled, dried in the SpeedVac, and then stored at -80ºC until further analysis.

For SDS-based processing, the embryos were mixed with 100 μl of SDS lysis buffer (4% SDS, 100 mM Tris-HCl, pH8.0), vortexed at 1,200 rpm for 5 min, and then sonicated under water bath for 5 min. The lysates were then mixed with final concentration of 10 mM TCEP plus 40 mM CAA, and boiled at 95ºC for 10 min. After cooling, the samples were processed following the E3filter procedure as reported previously.[2] In brief, the lysates were mixed with 80% ACN to induce protein aggregation, then transferred to E3filters (CDS Analytical, Oxford, PA). Next, the filters were washed 2-3 times with 80% ACN, spun at 1,000 x g for 1 min and the flow through was discarded. The samples were digested and desalted following the same procedure as described previously.[2]

#### LC-MS/MS analysis

The LC-MS/MS analysis was conducted with an Ultimate 3000 RSLCnano system coupled to an Orbitrap Eclipse mass spectrometer and FAIMS Pro Interface (Thermo Scientific). The peptides were resuspended in 20 μl of LC buffer A (0.1% formic acid in water), and half of it was loaded onto a trap column (PepMap100 C18, 300 µm × 2 mm, 5 µm; Thermo Scientific) followed by separation on an analytical column (PepMap100 C18, 50 cm × 75 µm i.d., 3 µm; Thermo Scientific) flowing at 250 nL/min. A linear LC gradient was applied from 1% to 25% mobile phase B over 125 min, followed by an increase to 32% mobile phase B over 10 min. The column was washed with 80% mobile phase B for 5 min, followed by equilibration with mobile phase A for 15 min. For the ion source settings, the spray voltage was set to 1.8 kV, funnel RF level at 50%, and heated capillary temperature at 275°C. The MS data were acquired in Orbitrap at 120K resolution, followed by MS/MS acquisition in data-independent mode following a protocol described previously.[3] The MS scan range (m/z) was set to 380-985, maximum injection time was 246ms, and normalized AGC target was 100%. For MS/MS acquisition, the isolation mode was Quadrupole, isolation window was 8 m/z, and window overlap was 1 m/z. The collision energy was 30%, Orbitrap resolution was 30K, AGC Target was 400K, and normalized AGC target was 800%. For FAIMS analysis, a 3-CV experiment (−40|-55|-75) was applied.

**Proteome quantitation and data analysis**

Mass spec data were processed using Spectronaut software (version 19.2)[4] and a library-free DIA analysis workflow with directDIA+ and the *X. tropicalis* protein database (UniProt 2024 release; 76,225 sequences). Briefly, the settings for Pulsar and library generation include: Trypsin/P as specific enzyme; peptide length from 7 to 52 amino acids; allowing 2 missed cleavages; toggle N-terminal M turned on; Carbamidomethyl on C as fixed modification; Oxidation on M and Acetyl at protein N-terminus as variable modifications; FDRs at PSM, peptide and protein level all set to 0.01; Quantity MS level set to MS2, and cross-run normalization turned on. Bioinformatics analyses including t-test, correlation, volcano plot and clustering analyses were performed using Perseus software (version 2.1.0) and Prism GraphPad (version 10 unless otherwise indicated. The MS raw files associated with this study have been deposited to the MassIVE server (https://massive.ucsd.edu/) with the dataset identifiers MSV000097784.

(4) Bruderer, R.; Bernhardt, O. M.; Gandhi, T.; Miladinović, S. M.; Cheng, L.-Y.; Messner, S.; Ehrenberger, T.; Zanotelli, V.; Butscheid, Y.; Escher, C.; Vitek, O.; Rinner, O.; Reiter, L. Extending the Limits of Quantitative Proteome Profiling with Data-Independent Acquisition and Application to Acetaminophen-Treated Three-Dimensional Liver

Microtissues. *Molecular & Cellular Proteomics* **2015**, *14* (5), 1400–1410. https://doi.org/10.1074/mcp.M114.044305.

### SUPPLEMENTARY FIGURES

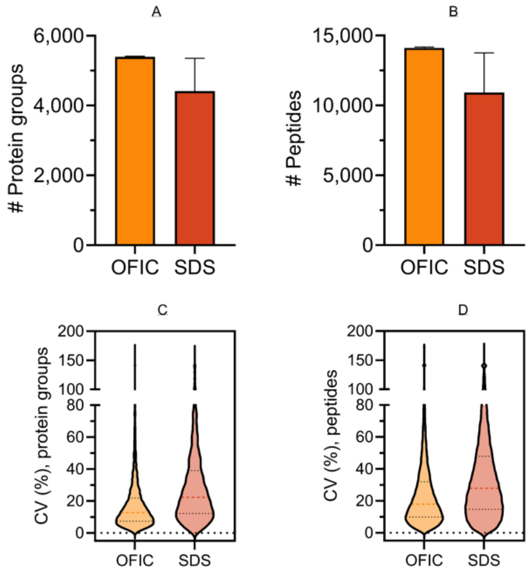


**Figure S1**. Comparison of on-filter in-cell (OFIC) digestion with SDS-based lysis method. (A-B) Protein and peptide identifications. Error bars represent three replicates. (C-D) Coefficient of variation of protein and peptide hits derived from the two methods.

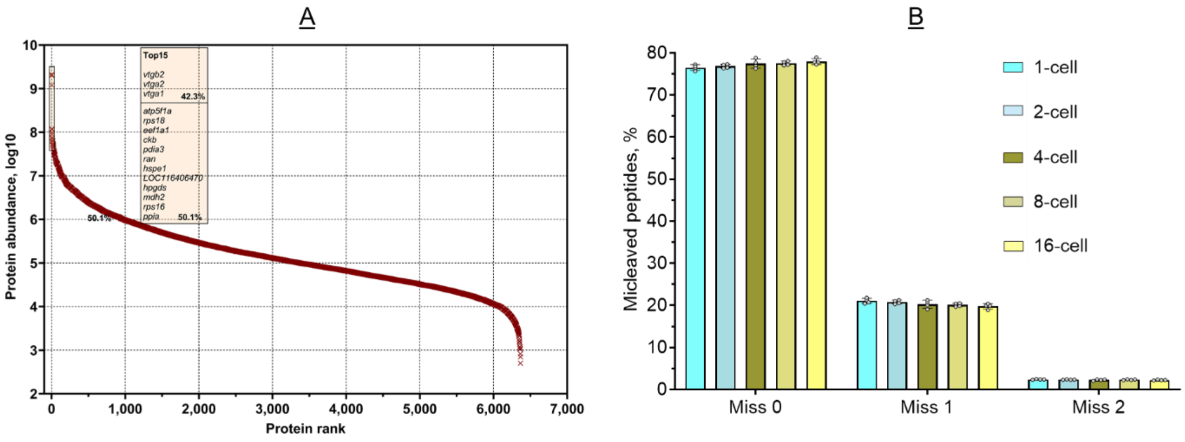


**Figure S2**. (A) The complete protein rank plot. The top 15 most abundant proteins were highlighted in the box. (B) Digestion efficiency of OFIC method calculated based on the percentage of the number of miscleaved peptides.

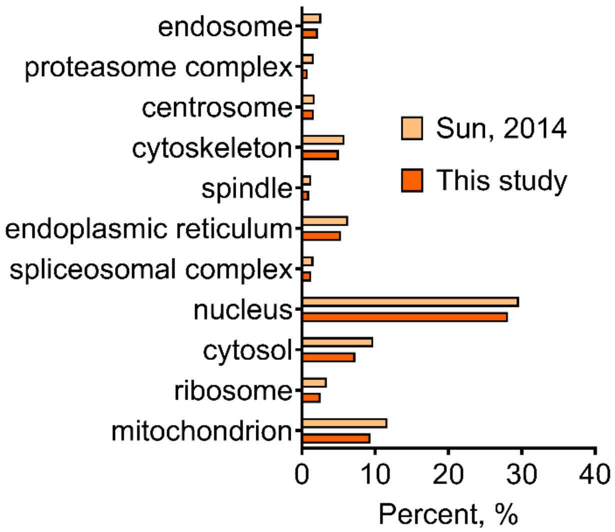


**Figure S3.** Comparison of *Xenopus* proteins from two studies using Gene Ontology cellular compartment analysis (GOTERM_CC_DIRECT). For this study, the total 6,375 *X. tropicalis* proteins were used for the analysis. For the study by Sun *et al.* [16], the total 4,065 *X. laevis* proteins were used.

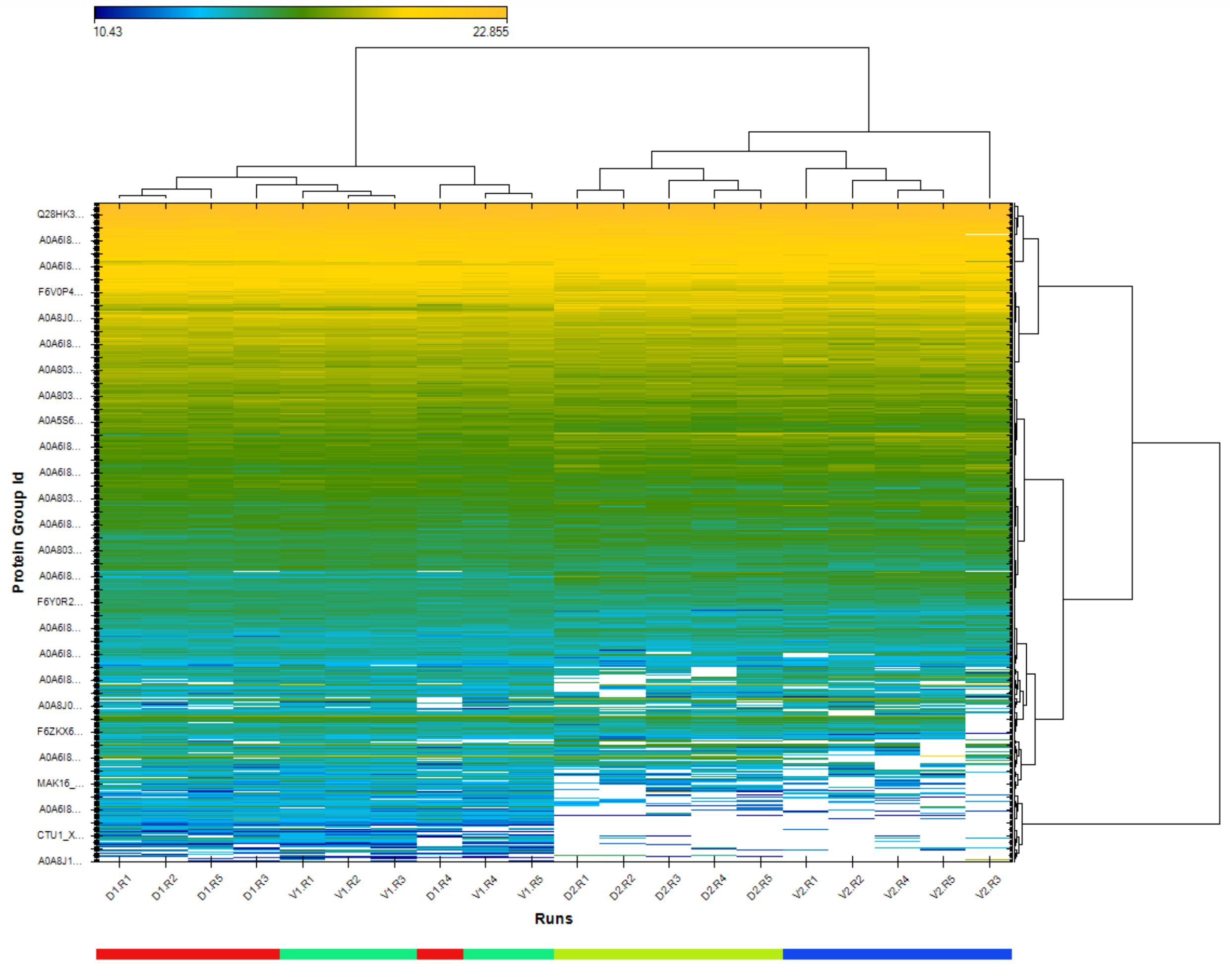


**Figure S4**. Unsupervised hieratical clustering analysis of the spatial proteomes of 8-Cell stage blastomeres. Five biological replicates were processed for each blastomere. White boxes indicate missing values.
